# Tubulin recycling limits cold tolerance

**DOI:** 10.1101/812867

**Authors:** Gabriella Li, Jeffrey K. Moore

## Abstract

Although cold temperatures have long been used to depolymerize microtubules, how temperature specifically affects the polymerization and depolymerization activities of tubulin proteins and how these lead to changes in microtubule networks in cells has not been established. We investigated these questions in budding yeast, an organism found in diverse environments and therefore predicted to exhibit dynamic microtubules across a broad range of temperatures. We measured the dynamics of GFP-labeled microtubules in living cells and found that lowering the temperature from 37°C to 10°C decreased the rates of both polymerization and depolymerization, decreased the amount of polymer assembled before catastrophes and decreased the frequency of microtubule emergence from nucleation sites. Lowering to 4°C caused rapid loss of almost all microtubule polymer. We provide evidence that these effects on microtubule dynamics may be explained in part by changes in the co-factor-dependent conformational dynamics of tubulin proteins. Ablation of tubulin-binding co-factors further sensitizes cells and their microtubules to low temperatures, and we highlight a specific role for TBCB/Alf1 in microtubule maintenance at low temperatures. Finally, we show that inhibiting the maturation cycle of tubulin by using a point mutant in β-tubulin confers hyper-stable microtubules at low temperatures, rescues the requirement for TBCB/Alf1, and improves the cold tolerance of the yeast. Together, these results reveal an unappreciated step in the tubulin cycle in cells and suggest that this step may be a key limiting factor in the thermal tolerance of organisms.

## Introduction

Microtubules are distinct from other cytoskeletal polymers in that they rapidly alternate between phases of polymerization and depolymerization, a non-equilibrium behavior known as ‘dynamic instability’ (Mitchison and Kirschner 1984). This behavior depends on a cycle of nucleotide-dependent conformational changes in the heterodimeric αβ-tubulin subunits that form microtubules (Alushin et al. 2014; Geyer et al. 2015; Hyman et al. 1995). Each heterodimer binds two molecules of GTP (Berry and Shelanski 1972; Weisenberg, Borisy, and Taylor 1968)). One GTP is sandwiched between α and β, and is neither exchanged nor hydrolyzed (Spiegelman, Penningroth, and Kirschner 1977). The other GTP binds to an exposed site on β, known as the exchangeable site or “E-site”. This nucleotide exchanges rapidly in solution and is hydrolyzed to GDP during microtubule polymerization (Carlier and Pantaloni 1981; Correia and Williams 1983; Purich and Kristofferson 1984). The binding of GTP to the E-site promotes tubulin polymerization and microtubule nucleation, so these rates exhibit a proportional dependence on the concentration of available GTP (Carlier, Didry, and Pantaloni 1987). Polymerization and nucleation rate also strongly depend on the concentration of tubulin (Olmsted et al. 1974; Walker et al. 1988). In the polymer state, tubulin hydrolyzes the E-site GTP and undergoes conformational changes that weaken binding to neighboring tubulins (Alushin et al. 2014; Carlier and Pantaloni 1981; Hyman et al. 1995). These conformational states have been described in great detail by recent cryo-EM studies, and the process of conformational changes is referred to as “maturation” (Alushin et al. 2014; Manka and Moores 2018; Zhang et al. 2015). GDP-bound tubulin dimers exhibit an approximately six-fold higher dissociation constant than GTP-bound tubulin, but can still contribute to microtubule assembly under *in vitro* conditions with sufficient concentrations of tubulin and Mg^2+^ (Carlier and Pantaloni 1978; Hamel et al. 1986). When microtubules switch from polymerization to depolymerization, an event known as catastrophe, mature GDP-bound tubulin is released from the lattice. In contrast to polymerization, depolymerization rates are concentration independent (Walker et al. 1988)

Although the assembly activity of tubulin and its relation to nucleotide status is well established, one important, but unanswered, question is whether tubulin visits intermediate, assembly-incompetent states between the free tubulin and microtubule states following microtubule disassembly. Previous studies have shown that cold temperatures can promote the formation of tubulin oligomers as a product of disassembly *in vitro*. As protofilaments peel away in a ‘ram’s horn’ shape, oligomers of curved tubulin are released from the microtubule (Bordas, Mandelkow, and Mandelkow 1983; Lange et al. 1988; Mandelkow, Mandelkow, and Milligan 1991). Increasing temperature causes these oligomers to dissociate into heterodimers (Bordas et al. 1983). Despite this evidence of temperature-dependent tubulin oligomers from *in vitro* experiments, what remains unknown is whether these structures occur *in vivo* and whether they represent points for regulating the pool of assembly-competent tubulin.

A potential mechanism for regulating the pool of assembly-competent tubulin would be to co-opt the tubulin biogenesis pathway. Tubulin biogenesis requires prefoldin and cytosolic chaperonin containing TCP-1 (CCT), to begin folding nascent α- and β-tubulin after translation ((Tian et al. 1996; Vainberg et al. 1998; Yaffe et al. 1992). However, unlike actin and other proteins that require CCT activity, tubulin also requires an additional set of tubulin-binding co-factors (TBCs) which bring together α- and β-tubulin subunits to form the assembly-competent heterodimer (Gao et al. 1993; Tian et al. 1997). In addition to their roles in tubulin biogenesis, TBCs also appear to regulate the activity of pre-formed heterodimers. *In vitro*, TBCC, TBCD, and TBCE form a complex that binds to pre-formed heterodimers and acts as a GTPase activator (GAP) for tubulin in the absence of microtubule polymerization (Nithianantham et al. 2015; Tian et al. 1999). Both TBCC and TBCE have each been shown to disassemble heterodimers *in vitro*, and overexpression of either in HeLa cells leads to microtubule loss (Bhamidipati, Lewis, and Cowan 2000). The dissociation of tubulin heterodimers by TBCE is enhanced greatly by the presence of TBCB, together forming a tri-partite complex of TBCE, TBCB and α-tubulin (Kortazar et al. 2007; Serna et al. 2015). These results demonstrate that TBCs can act on already formed tubulin heterodimers to alter nucleotide-binding status and interactions between tubulins. While it is unclear whether TBCs play a role in the transition of mature GDP-bound tubulin back to a GTP-bound assembly-competent heterodimer, it has been proposed that TBCs could provide a quality control mechanism to regulate the concentration of GTP-bound tubulin in cells (Tian and Cowan 2013).

The exquisite temperature sensitivity of microtubule dynamics provides a potential window for gaining insight into these questions. Early studies of cytoskeletal polymers found that microtubules are uniquely cold sensitive and quickly disappear upon incubation at low temperatures (Breton and Brown 1998; Tilney and Porter 1967; Weber, Pollack, and Bibring 1975; Weisenberg 1972). In fact, this property of tubulin is routinely used in protocols for purifying tubulin through cycles of temperature-induced polymerization and depolymerization (Borisy et al. 1975; Olmsted and Borisy 1973). The cold sensitivity of tubulin stands in contrast to F-actin polymers which persist after exposure to low temperatures (Breton and Brown 1998). The loss of microtubule polymer at low temperatures suggests a unique disruption of the normal tubulin cycle that may trap tubulin in an assembly-incompetent state.

In this study, we sought to determine how low temperature impacts the tubulin cycle to lead to the loss of microtubules *in vivo*. We used the budding yeast model, which has several major advantages for our study. Budding yeast have a simple microtubule network allowing for the measurement of single astral microtubules while possessing only one β-tubulin and two α-tubulin isotypes, thus forming a more homogeneous tubulin pool than that found in higher eukaryotes. Yeast are also found in diverse environments around the world, suggesting that yeast tubulin may exhibit a broader dynamic range of temperature response. We find that the rates of yeast microtubule polymerization and depolymerization are proportional to temperature and provide evidence that TBCs promote tubulin activity at low temperatures. We propose a model in which low temperatures trap tubulin in an assembly-incompetent state, and the pathway involved in tubulin biogenesis may help return it to assembly competence.

## Results

### Temperature dependence of microtubule dynamics *in vivo*

We began by testing the hypothesis that microtubules fail to assemble at low temperatures because the rate of microtubule polymerization may be more temperature dependent than the rate of depolymerization. This hypothesis predicts that the polymerization rate will decrease more sharply as temperature decreases, compared to the depolymerization rate. To test this prediction, we measured the lengths of single GFP-Tub1 labeled astral microtubules over time in cells incubated at different temperatures (Fig 1A, B). We controlled temperature using a stage-top microfluidic device, shifting from our standard temperature (30°C) to the target temperature for ~10 minutes before imaging (see Materials and Methods). In our experimental set up, 10°C was the lowest temperature that could be stably maintained during imaging. At the other end of our temperature range, we found similar phenotypes for 37°C and 39°C; but temperatures higher than 39°C severely inhibited microtubule assembly (data not shown). Within the range from 37°C to 10°C, we find that median astral microtubule length observed per cell over 10 minutes tended to decrease when the temperature was lowered (Fig 1C).

**Figure 1.**
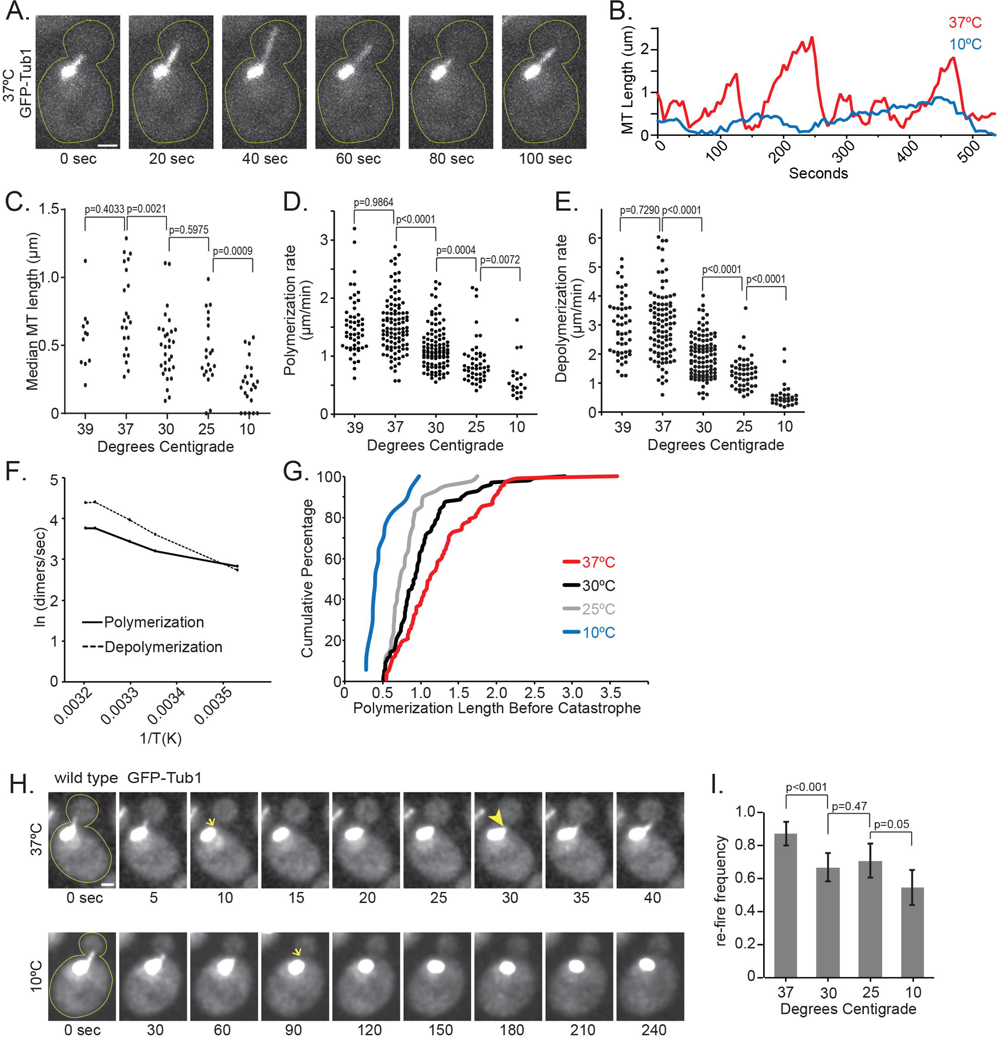
Temperature dependence of microtubule dynamics in vivo. A) Time-lapse image series of a wild-type cell expressing GFP-Tub1, showing the dynamics of a single astral microtubule at 37°C. Scale bar, 1um. B) Lifeplots of single astral microtubules at 10°C (blue) and 37°C (red). Microtubule lengths were measured at 5-second intervals. C) Median astral microtubule lengths from time-lapse imaging at indicated temperatures. Each dot represents the median length from 1 cell imaged for 10 minutes. For C-E, data were collected from at least three separate experiments; n= 23 cells at 10°C; n= 21 cells at 25°C; n= 30 cells at 30°C; n= 22 cells at 37°C; and n= 11 cells at 39°C. D) Polymerization rates of astral microtubules at indicated temperatures. Each dot represents a polymerization event. E) Depolymerization rates of astral microtubules at indicated temperatures. Each dot represents a depolymerization event. F) Arrhenius plot depicting the natural log of polymerization (solid line) and depolymerization (dashed line) rates extrapolated to dimers per second, as a function of the inverse temperature in Kelvin. G) Cumulative percentage of microtubule lengths at catastrophe at 10°C (Blue), 25°C (Grey), 30°C (Black), and 37°C (Red). H) Image series of wild-type cells with single dynamic astral microtubules labeled with GFP-Tub1 at 37°C (top) and 10°C (bottom). Arrows indicate when microtubule lengths drop below 0.213µm. Arrowhead indicates when a microtubule ‘re-fires’ in the cell at 37°C, and the length increases beyond 0.213µm. Scale bar, 1um. I) Re-fire frequencies calculated from cells imaged for 10 minutes at indicated temperatures. Values represent the sum of microtubules that re-fire within 20 seconds divided by the sum of microtubules that depolymerize to lengths less than 0.213µm, from at least 20 cells at each temperature. Error bars are standard error of proportion.

We used the full data set of microtubule length measurements over time to calculate different parameters of individual astral microtubule dynamics. As expected, polymerization rates of astral microtubules decrease as temperature decreases from 37°C to 10°C (1.589 ±0.1145 at 37°C, n=102; 1.151 ±0.0730 at 30°C, n=104; 0.9064 ±0.11745 at 25°C, n=47; 0.6226 ±0.15845 at 10°C, n=20; mean ±95%CI; Fig 1D). We also found that depolymerization rates decrease across this temperature range (3.013 ±0.2340 at 37°C, n=97; 1.944 ±0.1445 at 30°C, n=101; 1.360 ±0.1585 at 25°C, n=49; 0.5731 ±0.16255 at 10°C, n=29; mean ±95%CI; Fig 1E). The Arrhenius Plot in Figure 1F depicts the rates of change for polymerization and depolymerization across temperatures. As temperature decreases, the rate of depolymerization slows faster than the rate of polymerization, and these rates are predicted to intersect at ~13°C (Fig 1F). Based on these results, the decrease in microtubule polymerization rate at low temperatures (<13°C) would be expected to be offset by slower depolymerization. Therefore, the loss of microtubule polymer at low temperatures cannot be explained by changes in polymerization and depolymerization rates alone.

In addition to changes in polymerization and depolymerization, we identified other parameters that exhibit strong dependence on temperature. To determine how catastrophe is impacted by temperature change, we measured the length that a microtubule assembles before a catastrophe event and found that these assembly lengths decrease as temperature decreases (Fig 1G). In other words, at lower temperatures, less lattice is assembled before the microtubule undergoes catastrophe. We attempted a similar analysis to assess rescue events; however, the scarcity of rescue events at 10°C and 25°C does not permit a reliable measurement. At these lower temperatures, microtubules tend to depolymerize below our resolution limit (0.213 µm) and cannot be distinguished from the spindle pole bodies (SPBs). We also measured dynamicity, defined as the overall exchange of tubulin dimers at the microtubule end per second (Jordan et al. 1993). We found that dynamicity decreases significantly as the temperature lowers from 37°C to 10°C (51.07±5.82 at 37°C, n=22; 35.25±1.89 at 30°C, n=30; 25.98±2.47 at 25°C, n=21; 13.80±2.80 at 10°C, n=23; mean ±95%CI; Table 1).

**Table 1.** Microtubule dynamics across temperatures

| | Polymerization<br>($\mu\text{m}/\text{min}$) | Depolymerization<br>($\mu\text{m}/\text{min}$) | Polymerization<br>length before<br>catastrophe ($\mu\text{m}$) | Dynamicity<br>(subunits/sec) |
| --- | --- | --- | --- | --- |
| 10°C<br>(n=23) | 0.62 ± 0.16 | 0.57 ± 0.16 | 0.50 ± 0.21 | 13.80±2.80 |
| 25°C<br>(n=21) | 0.91 ± 0.12 | 1.36 ± 0.16 | 0.81 ± 0.17 | 25.98±2.47 |
| 30°C<br>(n=30) | 1.15 ± 0.07 | 1.94±0.15 | 1.01 ± 0.18 | 35.25±1.89 |
| 37°C<br>(n=22) | 1.59 ± 0.11 | 3.01±0.23 | 1.23 ± 0.21 | 51.07±5.82 |
| 39°C<br>(n=11) | 1.59±0.18 | 2.95±0.29 | | 49.48±5.41 |

We next examined how temperature impacts the formation of microtubules from nucleation sites at the SPBs. We identified astral microtubules that depolymerized to lengths below the resolution limit of our microscope (0.213µm) and measured the time elapsed before an astral microtubule re-emerged from the SPB (Fig 1H). At lower temperatures, we observed significant delays before re-emergence (Fig 1I; 10°C). Figure 1H depicts the frequency of astral microtubules that emerged from the SPB within 20 seconds of depolymerizing below our resolution limit; we termed this ‘re-fire frequency’. This analysis shows that low temperature inhibits astral microtubule emergence from the SPB (Fig 1I; S1). In fact, these data may underrepresent the impact of low temperature. We find that at 10°C, 7 of 22 cells failed to form new astral microtubules within the 10-minute timeframe of our analysis, compared to 0 of 30 cells at 30°C and 1 of 21 cells at 37°C. These results suggest that failure to initiate microtubule formation at nucleation sites may contribute to the loss of microtubule polymer at low temperature.

### αβ-tubulin levels are maintained at low temperature, despite loss of microtubules

Because microtubule polymerization rate depends on the concentration of αβ-tubulin heterodimers, we next tested whether the loss of microtubules at low temperature could be attributable to cold-induced destruction of αβ-tubulin heterodimers. Previous work in human neuronal cell lines and iPSC-derived neurons has shown that exposure to low temperatures causes a loss of microtubule polymer as well as a decrease in tubulin protein levels (Huff et al. 2010; Ou et al. 2018). To determine if budding yeast have the same response to low temperatures, we examined microtubules and αβ-tubulin protein levels in cells after prolonged exposure to low temperature. Cells grown to early log phase at 30°C and then shifted to 4°C lost all astral microtubules within 15 minutes, and this loss of polymer persisted after 24 hours at 4°C (Fig 2A, B). Nuclear microtubule loss was more gradual than astral microtubules, but after 60 minutes at 4°C, most cells lacked both astral and nuclear microtubules. In contrast to the loss of microtubule polymer, we found no change in tubulin protein levels, based on western blots for α-tubulin and β-tubulin in lysates from cells incubated at 30°C or 4°C for 24 hours (Fig 2C). As a second test of whether tubulin irreversibly loses activity during prolonged exposure to low temperature, we measured the kinetics of astral microtubule recovery after shifting cells from 4°C back to 30°C (Fig 2D). We found that astral microtubules recover quickly, on a time scale of seconds, when cells are shifted back to 30°C (Fig 2E). This indicates that αβ-tubulin can quickly recover polymerization activity when the temperature is raised. The rapid time scale also suggests that it is unlikely that recovery requires the biogenesis of a new pool of tubulin at 30°C, which occurs on the order of minutes (J. Moore, unpublished). These results suggest that low temperature inhibits microtubule assembly without depleting the levels of tubulin protein in the cell.

**Figure 2.**
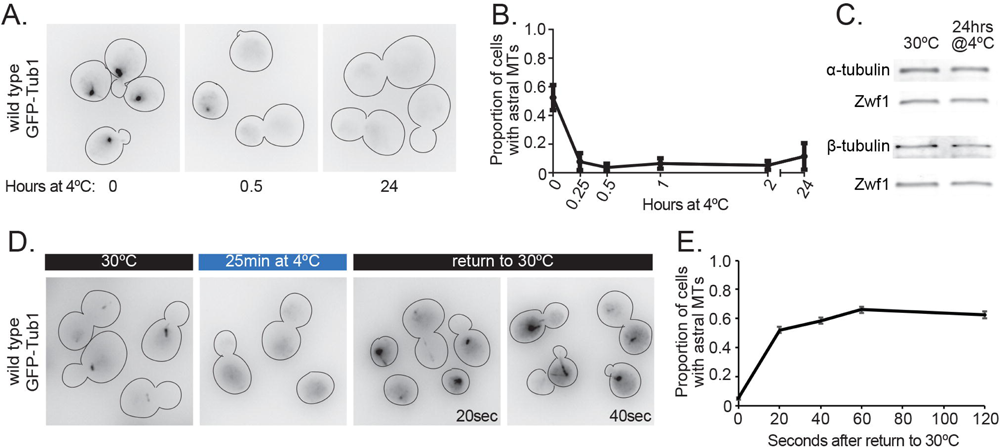
αβ-tubulin levels are maintained at low temperature, despite loss of microtubules. A) Representative field of wild-type cells expressing GFP-Tub1 shifted from 30°C to 4°C for 0, 0.5, and 24 hours. These images use an inverted lookup table to enhance contrast for the GFP-Tub1 signal. B) Proportion of wild-type cells with astral microtubules present after shifting to 4°C for indicated time. Values are mean ± SEM from at least three separate experiments, with at least 420 cells analyzed for each timepoint. C) Western blot of protein lysate from wild-type cells incubated at 30°C and at 4°C for 24hrs, probed for a-tubulin, -tubulin, and Zwf1. D) Representative field of wild-type cells expressing GFP-Tub1 incubated at 30°C, 25min after shifting to 4°C, and 20 and 40 seconds after shifting back to 30°C from 4°C. E) Proportion of wild-type cells with astral microtubules present after shifting from 4°C to 30°C for indicated time. Each data point represents a proportion calculated from data pooled from three separate experiments, with at least 482 cells analyzed for each timepoint. Error bars are standard error of proportion.

### Filamentous actin is not lost at low temperature

Low temperature is expected to generally slow biochemical reactions and have pervasive effects on cellular homeostasis that could indirectly inhibit microtubule assembly. To assess whether cytoskeletal dynamics are generally diminished at low temperatures, we examined effects on the actin cytoskeleton. We measured F-actin polymerization and depolymerization across temperatures using the well-established model of endocytic patch dynamics in budding yeast (Lin et al. 2010). We labeled F-actin using Lifeact-GFP and recorded time-lapse images on a spinning disk confocal microscope at 10°C and 37°C (Fig 3A). The lifetimes of actin patches are longer at 10°C, compared to 37°C (29.92 ±2.72 seconds at 10°C, n = 49; 7.08, ±0.786 seconds at 37°C, n = 30; p <0.0001; Fig 3A, B). Furthermore, the rates at which the intensity of Lifeact-GFP at individual patches increase and decrease, indicative of F-actin polymerization and depolymerization, are slower at 10°C compared to 37°C (Fig 3C). These slower rates are reminiscent of the effects we measured for microtubule dynamics. However, an important difference is that cells do not lose F-actin polymer at low temperatures. We shifted cells to 4°C for up to 24 hours and found that actin patches and cables are still present (Fig 3D). We conclude that actin dynamics slow at low temperatures, but unlike microtubules, F-actin is not lost.

**Figure 3.**
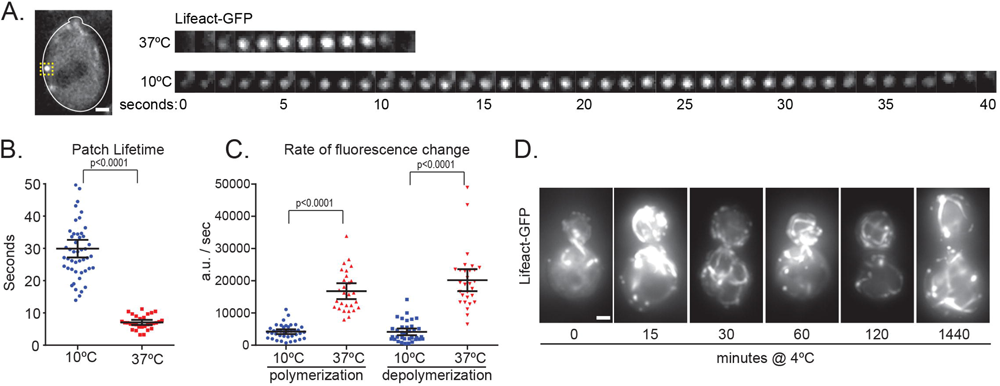
Filamentous actin is not lost at low temperature. A) Image of example wild-type cell expressing Lifeact-GFP with yellow box labeling an actin patch (left). Scale bar, 1um. Image series of zoomed in actin patch at 37°C (top) and 10°C (bottom) imaged at 1 second intervals (right). B) Actin patch lifetime measured by the full-width at ¾ maximum intensity of the actin patch curve at 10°C (blue) and 37°C (red). n= 49 cells at 10°c; n= 30 cells at 37°C; Values are mean with 95%C I. C) Rates of actin patch polymerization and depolymerization at 10°C (blue) and 37°C (red). n= 39 polymerization and 36 depolymerization events at 10°C; n= 28 polymerization and 29 depolymerization events at 37°C; Values are mean with 95%CI. D) Images series of wild-type cells expressing Lifeact-GFP shifted from 30°c to 4°c for the number of minutes indicated below. Scale bar, 1μm.

### Tubulin chaperones help recycle tubulin at low temperatures

We hypothesize that low temperature may be trapping tubulin in an assembly-incompetent state, such as the cold induced oligomers seen *in vitro* (Bordas et al. 1983; Lange et al. 1988; Mandelkow et al. 1991), and factors that bind tubulin could play a role in recycling a heterodimer back to an assembly-competent state. We sought to use cold sensitivity to identify cellular mechanisms for recycling assembly-incompetent tubulin by screening null alleles for a cold sensitive phenotype. We hypothesized that cells might co-opt tubulin biogenesis pathways to maintain the pool of assembly-competent tubulin (Fig 4A), and that this role may become more important at lower temperatures. To test this, we systematically generated all viable null mutants in the tubulin biogenesis pathway and screened for growth defects at low temperature (Fig 4B, C). We also screened mutations known to regulate microtubule dynamics and motor activity (Fig S2). To distinguish growth defects that are specific to cold sensitivity from those caused by general depletion of assembly-competent heterodimers, we also tested each mutant for sensitivity to the destabilizing drug benomyl (Fig 4B and C; S2). We found that null mutants in the prefoldin complex components *GIM3*, *GIM5*, and *YKE2* impair growth at 15°C, compared to wild-type controls, but also exhibit hyper-sensitivity to benomyl (Fig 4B). Although null mutants in most CCT genes are lethal in yeast, we did test a null mutant in *PLP1* (Lacefield and Solomon 2003), but found that this has little to no phenotype at low temperature or in the presence of benomyl (Fig 4B). Several null mutants in the TBC genes showed sensitivity to low temperature and benomyl; however, the null mutant in *ALF1*/*TBCB* showed increased cold sensitivity but only slightly increased benomyl sensitivity, compared to wild-type controls (Fig 4C). We therefore prioritized the *ALF1/TBCB* null mutant for further study. Our analysis also identified increased cold sensitivity for null mutants in the microtubule regulators Bik1/CLIP170, Bim1/EB1 and Cin8/kinesin-5 (Fig S2).

**Figure 4.**
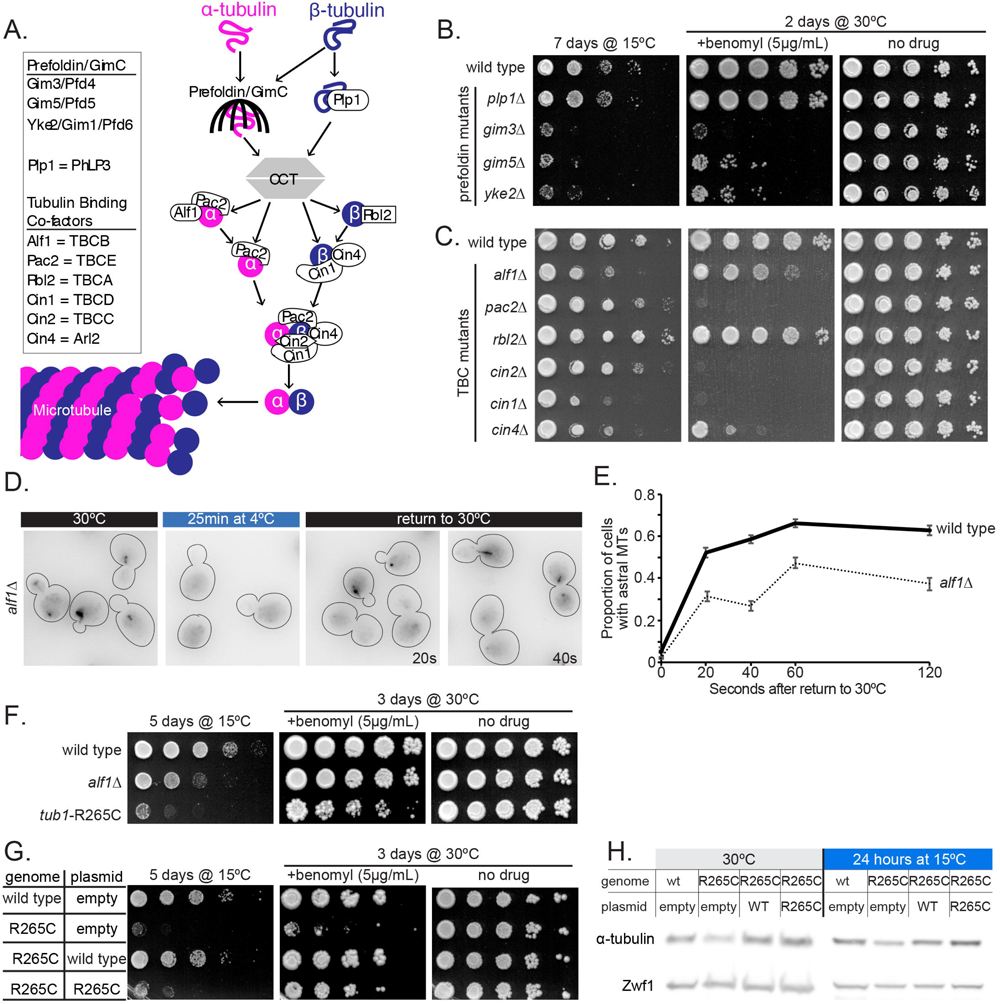
Tubulin chaperones help recycle tubulin at low temperatures. A) Model of the tubulin biogenesis pathway. (Adapted from Nithianantham et al., 2015; G Tian et al., 1997)C, D, and E. 8,C) Ten-fold dilution series of null strains for the prefoldin (8), CCT (8), and T8Cs (C) indicated at left were spotted to rich medium or rich medium supplemented with 5 µg/ml benomyl and grown at the indicated temperature for the indicated time. D) Representative field of alf1/1cells expressing GFP-Tub1 incubated at 30°C, 25min after shifting to 4°C, and 20 and 40 seconds after shifting back to 30°C from 4°C. E) Proportion of wild-type (solid line) and alf1/1(dotted line) with astral microtubules present after shifting from 4°C to 30°C for indicated time. Each data point represents a proportion calculated from data pooled from three separate experiments, with at least 266 cells analyzed for each timepoint. Error bars are standard error of proportion. F) Ten-fold dilution series of strain indicated at left were spotted to rich medium or rich medium supplemented with 5 µg/ml benomyl and grown at the indicated temperature for the indicated time. G) Ten-fold dilution series of strain carrying the plasmid indicated at left were spotted to rich medium, or rich medium supplemented with 5 µg/ml benomyl and grown at the indicated temperature for the indicated time. H) Western blot of protein lysate from wild-type or tub1-R265C cells carrying the indicated plasmid incubated at 30°C or 15°C for 24hrs, probed for a-tubulin and Zwf1.

To investigate how loss of *ALF1/TBCB* impacts tubulin activity at low temperature, we created an *alf1*∆ null mutant strain that expresses GFP-Tub1. We initially sought to measure microtubule dynamics in this strain; however, we found the GFP-Tub1 signal bleaches rapidly in cells lacking *ALF1/TBCB*. This does not appear to be due to lower expression of GFP-Tub1 in *alf1∆* mutants, based on western blotting (data not shown); and may therefore reflect a loss of assembly activity for the GFP-Tub1 fusion. Because these mutant cells are not amenable for time-lapse imaging, we instead used our assay to measure microtubule recovery upon raising the temperature from 4°C to 30°C. We find that the *alf1*Δ mutant slows the recovery of microtubules after exposure to low temperatures. When incubated at 4°C for 25 minutes before increasing the temperature back to 30°C, *alf1*Δ mutant cells exhibit slower recovery of astral microtubules than wild-type cells (Fig 4E). This slower recovery of astral microtubules in the *alf1*Δ mutant still happens on a time scale that is faster than would be expected for the biogenesis of new tubulin. Accordingly, this suggests that Alf1 promotes the recovery of tubulin assembly competence after cold induced depolymerization.

To further investigate a role for Alf1/TBCB in tubulin recycling, we used an α-tubulin mutation, R264C, that was originally identified through association with human brain malformations and subsequently shown to disrupt interaction with TBCB (Tian et al. 2008). We generated an analogous mutation at the R265 residue in the major yeast α-tubulin, *TUB1*. We find that *tub1*-R265C mutant cells exhibit a level of cold sensitivity that is similar to *alf1∆* null mutants (Fig 4F). However, *tub1*-R265C mutants are more sensitive to benomyl than *alf1∆* null mutants (Fig 4F). Providing cells with an additional copy of the *tub1*-R265C mutant allele rescues benomyl sensitivity, but not cold sensitivity (Fig 4G, H). This suggests the interaction between tubulin and the Alf1/TBCB becomes essential at low temperature. This supports our hypothesis that the tubulin biogenesis pathway plays a role in recycling tubulin to replenish the assembly-competent pool.

### Preventing tubulin maturation stabilizes microtubules at low temperatures

As a second test of our hypothesis, we predicted that tubulin mutants that persist in an immature, assembly-competent state could confer cold stability to microtubules. To test this prediction, we introduced a C354S substitution mutation into the β-tubulin gene, *TUB2* (Fig 5A). The *tub2*-C354S mutant has been previously shown to form hyper-stable microtubules *in vivo* and *in vitro*, and microtubules assembled from tub2-C354S heterodimers *in vitro* preferentially bind to the yeast EB protein, Bim1 (Geyer et al. 2015; Gupta et al. 2001, 2002). Based on these results, this mutation is thought to block heterodimer compaction associated with GTP hydrolysis, and therefore remain in a constitutively immature, assembly-competent state (Geyer et al. 2015). Consistent with previous studies, we find that cells expressing tub2-C354S exhibit persistent astral microtubules after shifting to 4°C (Fig 5B). These astral microtubules persist even after incubating mutant cells for 24 hours at 4°C (Fig 5C). In our microtubule dynamics assay, we find that tub2-C354S slows microtubule polymerization and depolymerization and exhibits fewer catastrophes and rescues than wild-type cells (Fig 5D). At low temperature (10°C), tub2-C354S microtubules exhibit little change in length over time, indicating that the cold stability of the mutant tubulin is not attributable to faster assembly rates (Fig 5D, (Gupta et al. 2002)). To test whether the cold-stability of tub2-C354S mutant tubulin could be explained by increased tubulin concentration, we measured αβ-tubulin levels via western blot and found that wild-type and *tub2*-C354S cells maintain similar amounts of tubulin across temperatures (Fig 5E). Thus, blocking tubulin maturation is sufficient to produce stable microtubules at low temperature.

**Figure 5.**
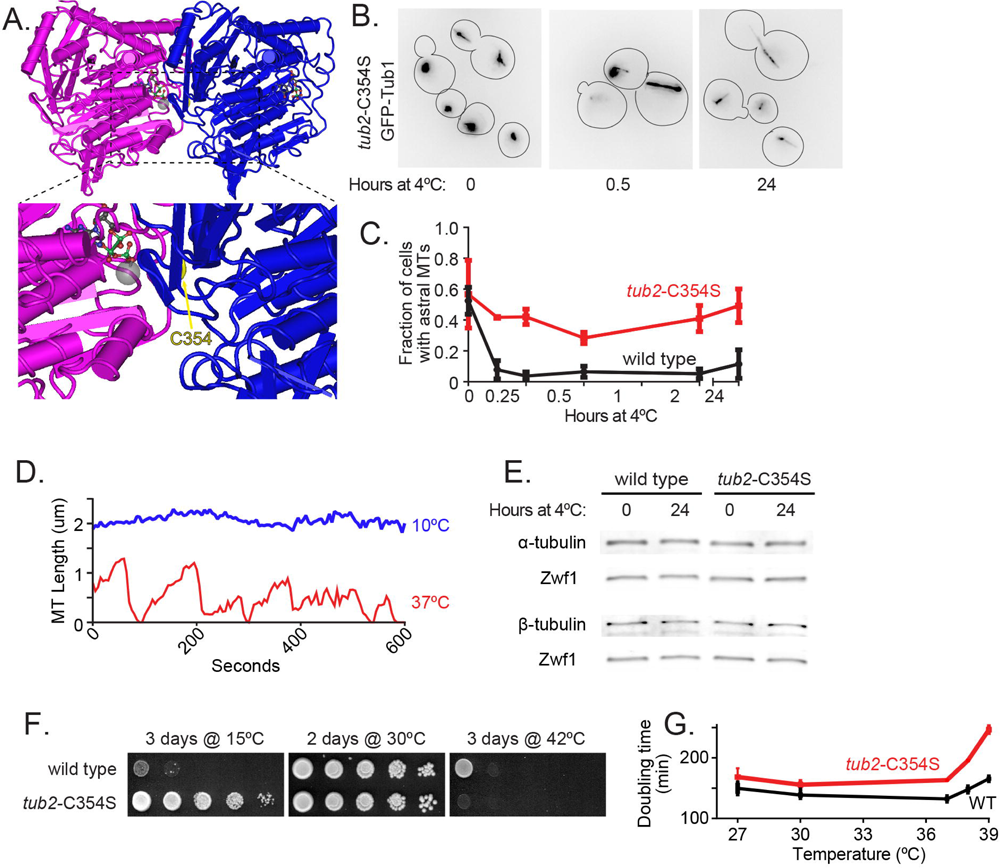
Preventing tubulin maturation stabilizes microtubules at low temperatures. A) Model of tubulin heterodimer with arrow pointing to residue C354 on -tubulin (PDB entry 5W3H; (Howes et al. 2017)). B) Representative field of tub2-C354S cells expressing GFP-Tub1 shifted from 30°C to 4°C for 0, 0.5, and 24 hours. These images use an inverted lookup table to enhance contrast for the GFP-Tub1 signal. C) Proportion of wild-type (black) and tub2-C354S (red) cells with astral microtubules present after shifting to 4°C for indicated time. Values are mean ± SEM from at least three separate experiments, with at least 420 cells analyzed for each timepoint. D) Lifeplots of a single tub2-C354S astral microtubule at 10°C (top) and 37°C (bottom). Each color represents one microtubule from three different cells. Microtubule lengths were measured at 5-second intervals. E) Western blot of protein lysate from wild-type (left) and tub2-C354S (right) cells incubated at 30°C and at 4°C for 24hrs, probed for a-tubulin, -tubulin, and Zwf1. F) Ten-fold dilution series of strains indicated at left were spotted to rich medium and grown at the temperature and amount of time indicated. G) Doubling times of wild type (black) and tub2-C354S (red) cells at the indicated temperature calculated from change in absorbance (OD600) over time. Values are mean ± SD from three separate experiments at each temperature.

Not only were tub2-C354S microtubules cold stable, but we also found that expressing tub2-C354S shifts the optimal temperature range for yeast cell growth. *tub2*-C354S mutant cells grow better at low temperatures than wild-type cells, but worse at high temperatures (Fig 5F, G; (Gupta et al. 2001)). To test if this cold tolerant growth was a result of tub2-C354S cells exhibiting more rapid cell division at lower temperatures, we used Spc110-tdTomato labeled strains to measure the rate at which cells released from α-factor synchronization at 15°C formed bipolar spindles (Fig S3B). We did not find a difference in the rate at which wild-type or *tub2*-C354S assemble spindles at 15°C (Fig S3C). As a positive control, we were able to detect a slowing of bipolar spindle formation in cells treated with benomyl to disrupt the microtubule network (Fig S3C). Thus, the improved growth of tub2-C354S cells at low temperature is not clearly attributable to improved spindle assembly; and the source of this phenotype remains unclear.

### *tub2*-C354S rescues the cold sensitivity of *alf1*Δ

Together, our data support a model in which low temperature traps mature tubulin in an assembly-incompetent state, and that the tubulin biogenesis pathway may help tubulin escape and return to the immature, assembly-competent state. Based on this model, we predict that the *tub2*-C354S mutation should rescue the requirement for *ALF1*/TBCB at low temperatures by keeping tubulin in an immature, assembly-competent state. To test this prediction, we generated double mutant cells combining *tub2*-C354S with *alf1*Δ and compared their growth at 15°C and in the presence of benomyl to that of the *alf1∆* single mutant. Consistent with our prediction, we find that the double mutant rescues the growth sensitivity of *alf1*Δ at low temperatures and sensitivity to benomyl (Fig 6A). As a second test, we directly assessed microtubule stability at low temperatures. We used a modified version of our cold-shift experiment – instead of incubating cells at 4°C, where wild-type cells lose all microtubule polymer, we used an intermediate temperature of 15°C and analyzed the portion of cells with remaining GFP-Tub1 labeled astral microtubules (Fig 6B). We find that *tub2*-C354S *alf1*∆ double mutants have a higher frequency of cells with astral microtubules than wild-type controls or the *alf1*Δ single mutant but decreased compared to the *tub2*-C354S single mutant after 120 minutes at 15°C (Fig 6C). We confirmed by western blot that all strains used in this experiment have similar tubulin concentrations, and that these concentrations are not altered by shifting to low temperature (Fig 6D). Therefore, the *tub2*-C354S mutation rescues the loss of *ALF1/TBCB* by restoring the quality of the tubulin pool, not the quantity. These results add support to our model that Alf1/TBCB promotes the return of tubulin to an assembly-competent state.

**Figure 6.**
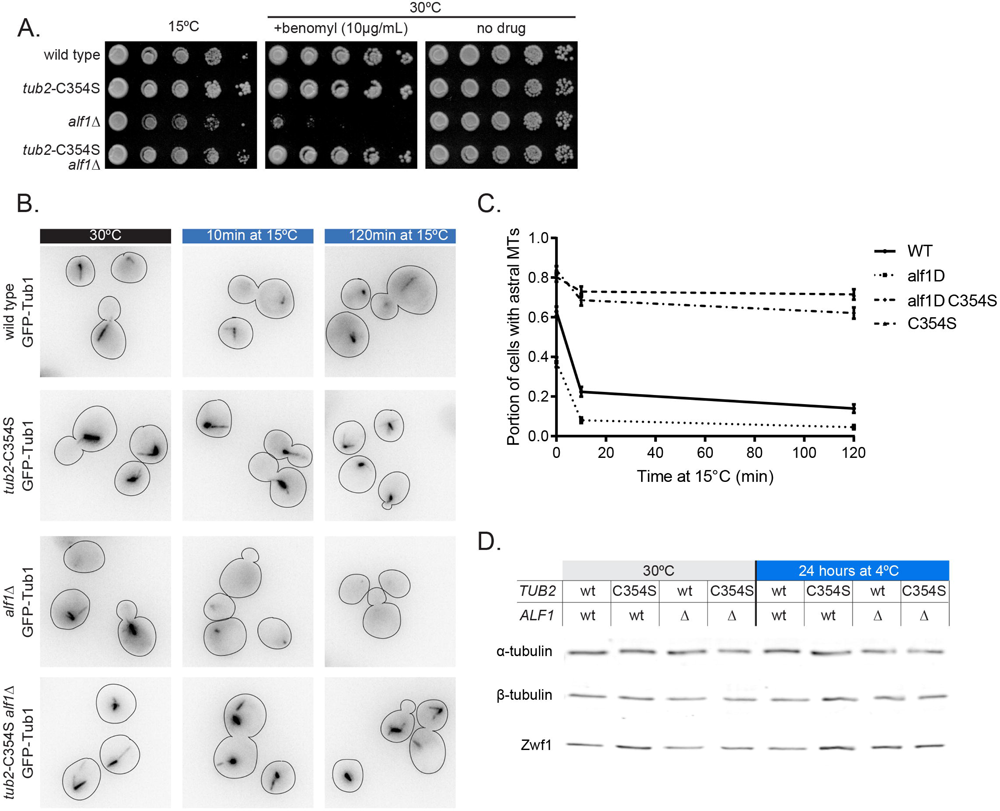
tub2-C354S rescues the cold sensitivity of alf1Δ. A) Tenfold dilution series of strains indicated at left were spotted to rich medium or rich medium supplemented with 10 µg/ml benomyl and grown at the indicated temperature for the indicated time. B) Representative field of wild-type, alf1/1, tub2-C354S, and alf1/1with tub2-C354S cells expressing GFP-Tub1 after 20min at 30°C, 10min after shifting to 15°C, and 120min after shifting to 15°C. C) Proportion of wild-type (solid line), alf1/1(dotted line), tub2-C354S (dashed line), and alf1/1with tub2-C354S (dot-dashed line) cells with astral microtubules present after shifting from 30°C to 15°C for indicated time. Each data point represents a proportion calculated from data pooled from three separate experiments, with at least 293 cells analyzed for each timepoint. Error bars are standard error of proportion. D) Western blot of protein lysate from wild-type and tub2-C354S cells with or without Alf1 incubated at 30°C and at 4°C for 24hrs, probed for a-tubulin, αβ-tubulin, and Zwf1.

## Discussion

The cold sensitivity of microtubules has been known for many years. In early studies, cold sensitivity distinguished microtubules from other cytoskeletal filaments, and it has since been used as a tool to selectively disrupt microtubule networks in cells and to purify tubulin from cells through cycles of temperature-induced depolymerization and polymerization (Inoué 2010; Olmsted et al. 1974; Tilney and Porter 1967). Although cold sensitivity is well known, why cells lose microtubules at low temperatures has not been precisely defined. The goals of this study are to define the mechanistic basis for microtubule loss at low temperatures, and then to use this as a tool to gain broader insight into how cells regulate microtubule dynamics.

An important question is whether microtubule loss at low temperature is attributable to direct effects on tubulin activity, or indirect consequences of temperature-induced changes in cell physiology. While we cannot completely rule out the impact of other cold-induced changes in cell physiology, several lines of evidence indicate that microtubule loss results from direct effects on tubulin activity. It is well established that the activity of purified tubulin exhibits strong dependence on temperature. Microtubules reconstituted *in vitro* from purified mammalian tubulin are destabilized by low temperature and exhibit unique, curled protofilament architectures that suggest temperature-dependent changes in inter- and/or intra-heterodimer interactions (Mandelkow et al. 1991). Interestingly, tubulin purified from psychrophilic organisms forms microtubules that remain stable at low temperature, consistent with the notion that interspecies differences in the tubulin proteins may determine cold sensitivity (Himes and Detrich 1989). Thus, temperature directly and potently impacts the tubulin protein in a way that alters its assembly activity. By comparison, F-actin reconstituted *in vitro* from purified mammalian actin is not destabilized by low temperature ((Breton and Brown 1998; Weber et al. 1975). We also find F-actin in yeast cells is not lost at low temperatures (Figure 3). Although the rates of F-actin polymerization and depolymerization do slow when temperature decreases to 10°C, there is no net loss of F-actin, even after prolonged exposure to 4°C (Figures 3C). Thus, polymer loss at low temperature is a unique feature of tubulin and likely due to temperature-dependent effects on protein activity.

In seeking to define the mechanistic basis for microtubule loss at low temperatures, we first considered a simple kinetic model in which changes in four parameters of microtubule dynamics – polymerization rate, depolymerization rate, catastrophe frequency, and rescue frequency – would be sufficient to explain microtubule loss at low temperatures. We find that both polymerization and depolymerization rates decrease as the temperature decreases from 37°C and 10°C (Figure 1; Table 1) consistent with what has been observed *in vitro* (Fygenson, Braun, and Libchaber 1994). Surprisingly, the effect of temperature on depolymerization rate is more pronounced than what we observed for polymerization rate. The Arrhenius plot in Figure 1F shows these rates intersecting at ~13°C, and at lower temperatures the polymerization rate is faster than the depolymerization rate. Although polymerization and depolymerization are clearly impacted by temperature change, we conclude that these effects alone are not sufficient to explain the loss of microtubule polymer at low temperatures. We did not observe enough rescue events at 10 °C and 25 °C to make reliable measurements; however, we did find that catastrophe frequency increases at low temperature. Since polymerization slows as temperature decreases, we measured catastrophe as a function of microtubule length assembled, rather than as a function of time. We find that this length value decreases as the temperature lowers (Figure 1; Table 1). Thus, at low temperatures, less microtubule lattice is assembled before catastrophe occurs. We propose two models that could explain how low temperature promotes catastrophe. In the first model, the slower rate of tubulin addition to the plus end at low temperatures may not outpace GTP hydrolysis. Accordingly, plus ends may have shorter GTP caps at low temperatures. In the second model, low temperature may destabilize the plus end by inhibiting lateral interactions between protofilaments. This model is inspired by recent observations that assembling plus ends contain protofilament extensions that must zipper together to form a complete lattice (McIntosh et al. 2018), and older evidence that low temperature induces protofilaments to adopt an outwardly curled configuration that prevents lateral interactions (Mandelkow et al. 1991). We speculate that the increase in catastrophes may explain why microtubule polymer is lost at lower temperatures.

In addition, our results also suggest that low temperature inhibits microtubule nucleation activity. In this model, microtubules are lost at low temperatures because they depolymerize completely and then tubulin fails to nucleate new microtubules. Consistent with this, we find that low temperature inhibits the emergence of microtubules from SPBs (Figure 2A and B). Since SPBs contain Ɣ-tubulin small complexes (Ɣ-TuSCs) that nucleate microtubules, this effect could be attributable to cold sensitivity of Ɣ-TuSC activity or the intrinsic nucleation activity of αβ-tubulin heterodimers (Sobel and Snyder 1995; Zheng et al. 1995). Distinguishing between these possibilities will require further study.

We speculate that the formation of assembly-incompetent tubulin may be a pervasive feature of microtubule dynamics that is exacerbated by, rather than unique to, low temperature. Numerous *in vitro* reconstitution studies have shown that depolymerized tubulin is intrinsically capable of cycling back to an assembly-competent, GTP-bound state when provided with optimal conditions. These conditions that include warm temperatures, high concentrations of free GTP (1mM), 80mM PIPES, high Mg^2+^ (≥ 1mM), and low Ca^2+^(Borisy et al. 1975; Lee and Timasheff 1977; Weisenberg 1972). How this transition from mature GDP-tubulin to assembly-competent, GTP-tubulin is achieved in a complex *in vivo* cellular environment is an important question. Conditions in cells differ from *in vitro* reconstitution experiments in ways that are likely to impact tubulin activity. For example, consider budding yeast in which the total intracellular concentration of GTP is reported to be 1.5mM (Breton et al. 2008) with many proteins competing with tubulin for binding to this nucleotide pool. Intracellular Mg^2+^ ranges between 500uM and 1mM (Romani and Scarpa 1992), and most of these cations are complexed with proteins and nucleic acids (Cyert and Philpott 2013). Available Mg^2+^ *in vivo* is therefore likely below the concentrations required for microtubule dynamics *in vitro*. Finally, many eukaryotic cells maintain nM concentrations of Ca^2+^ in the cytoplasm, and budding yeast can survive with cytosolic Ca^2+^ concentrations up to 2mM (Halachmi and Eilam 1989; Kováč 1985); however, *in vitro* microtubule dynamics assays typically contain no Ca^2+^(Gell et al. 2010). Thus, the conditions that support dynamic tubulin assembly *in vitro* are unlikely to represent the complexity of tubulin regulation *in vivo*. Extrinsic factors are likely to play critical roles in regulating tubulin activity in the cellular environment.

Our results also reveal an additional effect of temperature on tubulin where low temperature causes tubulin to accumulate in an assembly-incompetent state that is the product of microtubule disassembly. Our confirmation of the previous finding that the C354S mutation in β-tubulin maintains microtubules at low temperature indicates that a full maturation cycle is required for tubulin to visit the assembly-incompetent state (Figure 5;(Gupta et al. 2001, 2002)). The C354S mutation is thought to disrupt the tubulin cycle and maintain an ‘immature’ state by uncoupling GTP hydrolysis from conformational changes (Geyer et al. 2015). Accordingly, C354S mutant tubulin may avoid the assembly-incompetent state by not visiting the mature state of the tubulin cycle. Importantly, we found that total tubulin concentration in yeast cells, for both wild-type and C354S mutants, does not change when temperature is decreased as low as 4°C (Figure 2E and 5D). This stands in contrast to the decrease in tubulin concentration during exposure to low temperatures that has been previously reported in cultured human neurons (Huff et al. 2010; Ou et al. 2018). Based on this evidence, we propose that when tubulin depolymerizes at low temperature it becomes trapped in a conformationally matured, assembly-incompetent state that is incompatible with microtubule polymerization. This model could therefore lead to microtubule loss by sequestering the pool of assembly-competent tubulin contributing to slower polymerization rates observed at low temperatures.

Our results indicate that TBCs, particularly TBCB/Alf1, may play an important role in recycling tubulin from an assembly-incompetent state to maintain the pool of assembly-competent tubulin in the cell. Interestingly, previous studies found that a complex of TBCB and TBCE dissolves heterodimers and forms a tri-partite complex with α-tubulin (Kortazar et al. 2007; Serna et al. 2015). The relevance of this activity in cells has not been established, but it can be speculated that this disassembly activity could be involved in heterodimer quality control or oligomer dissociation. We find that disrupting individual TBCs and other tubulin biogenesis factors further sensitizes cells to low temperatures and delays the recovery of microtubule polymerization after cold shock (Figure 4). Blocking α-tubulin from interacting with TBCB/Alf1 also sensitizes cells to cold temperatures (Figure 4). We reason that at low temperatures, the role of TBCs in maintaining the tubulin pool may become more important to help tubulin escape the assembly-incompetent state. How TBCs might inhibit or resolve this tubulin state, and whether this role extends to other stress conditions, are important questions for further study.

Finally, our work highlights a critical role for tubulin in determining the fitness of organisms at low temperature. We propose that shifting tubulin from a predominantly microtubule polymer state to cytoplasmic oligomers could impair homeostasis at low temperatures. Consistent with this, it is striking that a single point mutation, C354S, in β-tubulin confers cold stability to microtubules and also shifts the optimal growth temperature range of yeast cells, with mutant cells exhibiting improved growth at low temperatures but greater sensitivity to high temperatures, compared to wild-type cells (Figure 5). We find that the cold sensitivity of wild-type cells is not primarily due to mitotic spindle defects, since C354S mutants do not exhibit faster spindle assembly at low temperatures and *mad2*Δ mutant cells that lack the spindle assembly checkpoint show a similar growth rate at low temperatures to wild-type controls (Figure S3). It is possible that the increasing pool of tubulin oligomers at low temperature triggers a secondary stress response that is responsible for the slowing of cell proliferation. This suggests that microtubule stability is an integral factor in determining the permissive growth temperatures of an organism.

## Materials and Methods

### Yeast Strains, manipulations, and plasmid construction

General yeast manipulation, media and transformation were performed by standard methods(Amberg, Burke, and Strathern 2000). A detailed list of strains and plasmids is provided in Tables S1 and S2. GFP-Tub1 fusions were integrated at the *LEU2* locus and expressed ectopically, in addition to the native *TUB1* (Song and Lee 2001). Spc110-tdTomato was generated using conventional methods and expressed from the genomic locus (Sheff and Thorn 2004). The tub2-C354S mutation to *TUB2* was made at the native chromosomal locus as previously described (Aiken et al. 2014) and confirmed by sequencing. Deletion mutants were generated by conventional methods (Petracek and Longtine 2002). A fragment of the TUB1 gene including the open reading frame (with intron), 992 base pairs of 5’ UTR, and 487 base pairs of 3’ UTR was amplified from the genomic DNA by PCR. This fragment was cloned into the plasmid pRS314 (Sikorski and Hieter 1989) using sticky end cloning with NotI and KpnI sites, and confirmed by sequencing to yield plasmid pJM267. QuikChange Lightning Site-Directed Mutagenesis Kit (Agilent Technologies; Santa Clara, CA) was used to introduce the tub1-R265C mutation to pJM267 (Aiken et al. 2019).

### Microtubule Dynamics in Living Cells

Images were collected on a Nikon Ti-E microscope equipped with a 1.45 NA 100× CFI Plan Apo objective, piezo electric stage (Physik Instrumente; Auburn, MA), spinning disk confocal scanner unit (CSU10; Yokogawa), 488-nm laser (Agilent Technologies; Santa Clara, CA), and an EMCCD camera (iXon Ultra 897; Andor Technology; Belfast, UK) using NIS Elements software (Nikon). Cells were grown asynchronously to early log-phase in non-fluorescent media and adhered to coverslips coated with concanavalin A (Fees, Estrem, and Moore 2017). Imaging was performed at the indicated temperature using the CherryTemp temperature controller system (CherryBiotech; Rennes, France).

Microtubule dynamics were analyzed by measuring the lengths of astral microtubules labeled with GFP-Tub1 at 5-s intervals for 10 min. A 6.0 µm stack with a step size of 0.45 µm was taken at each timepoint and analyzed as a 2D maximum intensity projection (ImageJ; Wayne Rasband, NIH). All analyses were conducted in pre-anaphase cells. Assembly and disassembly events were defined as at least three contiguous data points that produced a length change ≥0.5 µm for 25°C, 30°C, and 37°C with a coefficient of determination ≥0.8. For experiments at 10°C, we adjusted the threshold for length change to ≥0.25 µm. Microtubule dynamicity was calculated by the total change in length (growing and shrinking) divided by the change in time and expressed in tubulin subunits changed per second (Jordan et al. 1993). The length of polymerization before a catastrophe event was calculated by determining the total length of polymerization before a switch to depolymerization. Median microtubule length was determined from time-lapse imaging of individual astral microtubules over 10 min. Dynamics measurements for individual microtubules were pooled for each temperature and then compared with pooled data for different temperatures. Student’s t test was used to assess whether the mean values for different datasets were significantly different (Fees et al. 2017). Arrhenius plot was generated by converting the polymerization and depolymerization rates from µm/min to heterodimers/sec, assuming that 1µm of yeast microtubule contains 1625 tubulin heterodimers.

Re-fire frequency was calculated as the proportion of astral microtubules that depolymerize to a length below our level of detection (< 0.213 µm) and then polymerized to a measurable microtubule length (> 0.213 µm) within 20 seconds (4 frames in our analysis), divided by the total number of astral microtubules that depolymerize to a length below our level of detection. At least 21 astral microtubules were analyzed for each temperature.

### Microtubule Loss at 4°C

Cells expressing GFP-Tub1 were grown overnight in rich liquid media in a shaking incubator at 30°C for ~16 hours, diluted 1:150, and grown to early log phase in fresh media. The cultures were then shifted to a shaking incubator at 4°C for the indicated amount of time. Cells were then fixed in 3.7% Formaldehyde, 0.1 M KPO_4_ and incubated at 4°C for 3 minutes. Cells were then pelleted, the supernatant was removed, and the cells were suspended in quencher solution (0.1% Triton-X, 0.1 M KPO_4_, 10 mM Ethanolamine). The cells were pelleted again, supernatant decanted and washed twice in 0.1 M KPO_4_. The fixed cells were loaded into slide chambers coated with concanavalin A, washed with 0.1 M KPO_4_ and the chambers were sealed with VALAP (Vaseline, lanolin and paraffin at 1:1:1) (Fees and Moore 2018).

Images were collected on a Nikon Ti-E wide field microscope equipped with a 1.49 NA 100× CFI160 Apochromat objective, and an ORCA-Flash 4.0 LT sCMOS camera (Hammamatsu Photonics, Japan) using NIS Elements software (Nikon, Minato, Tokyo, Japan). Microtubules labeled with GFP-Tub1 were imaged in Z-series consisting of a 7 µm range at 0.5 µm steps, and a DIC image was taken at the equatorial plane.

Cells were segmented using a custom ImageJ macro previously described (Fees et al. 2016). The data were blinded for analysis, and the segmented cells were visually scored for the presence of astral microtubules.

### Microtubule Recovery Assay

Cells expressing GFP-Tub1 were grown in rich liquid media in a shaking incubator at 30°C for ~16 hours, then diluted 1:150 and grown to early log phase in fresh media. 100 µL of each culture was transferred to 200 µL PCR tubes and placed in a Bio-Rad T100 Thermal Cycler (Hercules, CA). The cells were incubated at 4°C for 35 min and warmed to 30°C for the indicated amount of time before being fixed, imaged and analyzed for the presence of astral microtubules, as described above.

### Microtubule Loss at 15°C

Cells expressing GFP-Tub1 were grown in rich liquid media in a shaking incubator at 30°C for ~16 hours, then diluted 1:150 and grown to early log phase in fresh media. 100 µL of each culture was transferred to 200 µL PCR tubes and placed in a Bio-Rad T100 Thermal Cycler (Hercules, CA). The cells were incubated at 30°C for 20 min and cooled to 15°C for the indicated amount of time before being fixed, imaged and analyzed for the presence of astral microtubules, as described above.

### Actin Patch Dynamics in Living Cells

Images were collected on a Nikon Ti-E microscope described above. Actin patch lifetime and dynamics were analyzed by measuring the accumulation and decay of Lifeact-GFP in single z-plane images acquired every 1-s for 3 min. A 0.852 um X 0.852 um region of interest was used to measure total fluorescence of each patch. The actin patch lifetime was defined as the amount of time the patch was ≥ ¾ of its maximum intensity by measuring the full width at ¾ max of the fluorescence over time. Actin patch polymerization dynamics were determined by measuring the slope of fluorescence intensity versus time from before the patch appears to the maximum intensity, and depolymerization measured by the slope from maximum intensity to patch disappearance with a coefficient of determination ≥0.8.

### Western Blot

To compare tubulin protein levels before and after cold shift, cells were first grown to log phase at 30°C in 5 ml rich liquid media or selective drop out media for strains containing a plasmid, and 1 ml was removed for lysate preparation. The remaining cell culture was transferred to a platform shaking at 250rpms at 4°C for 24 hours, and then 1 ml was removed for lysate preparation. Samples were pelleted and then resuspended in 2 M lithium acetate for 5 minutes. The cells were then pelleted again and resuspended in 0.4 M NaOH for 5 minutes while on ice. Samples were then pelleted, resuspended in 70 µL of 2.5X Laemmli buffer and boiled for 5 minutes. The total protein concentration of the clarified lysate was determined by Pierce 660 nm protein assay with the Ionic Detergent Compatibility Reagent (Cat.1861426 and 22663, Rockford, IL), and ~5 ug of total protein lysate was loaded in each lane. Samples were run on 12% SDS PAGE, transferred to PVDF membrane, and blocked for 1 hour at room temperature. Membranes were probed with mouse-anti-α-tubulin (4A1; at 1:100; (Piperno and Fuller 1985)), mouse-anti-β-tubulin (E7; at 1:100; Developmental Hybridoma Studies Bank, University of Iowa) and rabbit-anti-Zwf1 (Sigma A9521; at 1:10,000), followed by goat-anti-mouse-680 (LI-COR 926-68070, Superior, NE; at 1:15000) and goat-anti-rabbit-800 (LI-COR 926-32211; at 1:15000), and imaged on an Odyssey Imager (LI-COR Biosciences). Band intensities were quantified using ImageJ.

### Liquid Growth Assay

Cells were grown to saturation in rich liquid media, diluted 1:1000 into fresh media, and transferred to a 96-well plate at a volume of 200ul/well. Absorbance values at 600 nm were measured at 5-minute intervals over 21 hours at specified temperature with orbital shaking using a Cytation 3 Plate Reader (Biotek; Winooski, VT). Doubling time was calculated by determining the OD600 values to an exponential curve as using a custom Matlab code as described previously (Fees and Moore 2018). Doubling times are based on data from 3 separate experiments, and at least 3 independent isolates for each genotype.

### Solid Growth Assay

Cells were grown in rich liquid media or selective drop out media for strains carrying a plasmid to saturation at 30°C, and a 10-fold dilution series of each culture was spotted to either rich media plates or rich media plates supplemented with 5 or 10 µg/ml benomyl. Plates were grown at the indicated temperature for the indicated amount of time.

### Synchronized Cell Cycle experiment

Cells were grown overnight in rich liquid media in a shaking incubator at 30°C to early log phase. Cells were pelleted, resuspended in fresh media with 5ug/ml α-factor, returned to the shaking incubator at 30°C for 1.5 hours, and then an additional 5ug/ml of α-factor was added for an additional 1.5 hours at 30°C. Cells were then washed three times with sterile water and resuspended in fresh media with 50ug/ml s. gris protease added for synchronous release through START. Wild-type cells were treated with either 1% DMSO or 5ug/ml benomyl in DMSO as indicated. The cells were then incubated at 15°C while shaking and fixed at the indicated timepoint as described above. Microtubule labeled with GFP-Tub1 and spindle pole bodies labeled with Spc110-tdtomato were imaged as described above. Cells were segmented using a custom ImageJ macro previously described (Fees et al. 2016). The data was blinded for analysis, and the segmented cells were visually scored for cell cycle stage.

## Acknowledgements

We thank members of the Moore lab and Dan Sackett (NIH NICHD) for helpful advice and discussions. This work was supported by the National Science Foundation CAREER Award 1651841 to J.K. Moore. G. Li was supported by the Pre-doctoral Training Program in Molecular Biology, NIH-T32-GM008730, and Bolie Scholar Award from the Graduate Program in Molecular Biology.

---

**Tables S1.** Plasmids used in this study

| Plasmid ID | Plasmid Name | Source |
| --- | --- | --- |
| pJM0101 | pRS415 *P<sub>CYC1</sub>*-Lifeact-GFP *LEU2 CEN6* | Lin et al. 2010 |
| pJM0072 | pRS314 *TRP1 CEN6* | Sikorski, R. S. and Hieter, P. 1989 |
| pJM0276 | pRS314 *TUB1 TRP1 CEN6* | Aiken et al. 2019 |
| pJM0296 | pRS314 *tub1-R265C TRP1 CEN6* | Aiken et al. 2019 |

**Tables S2.**
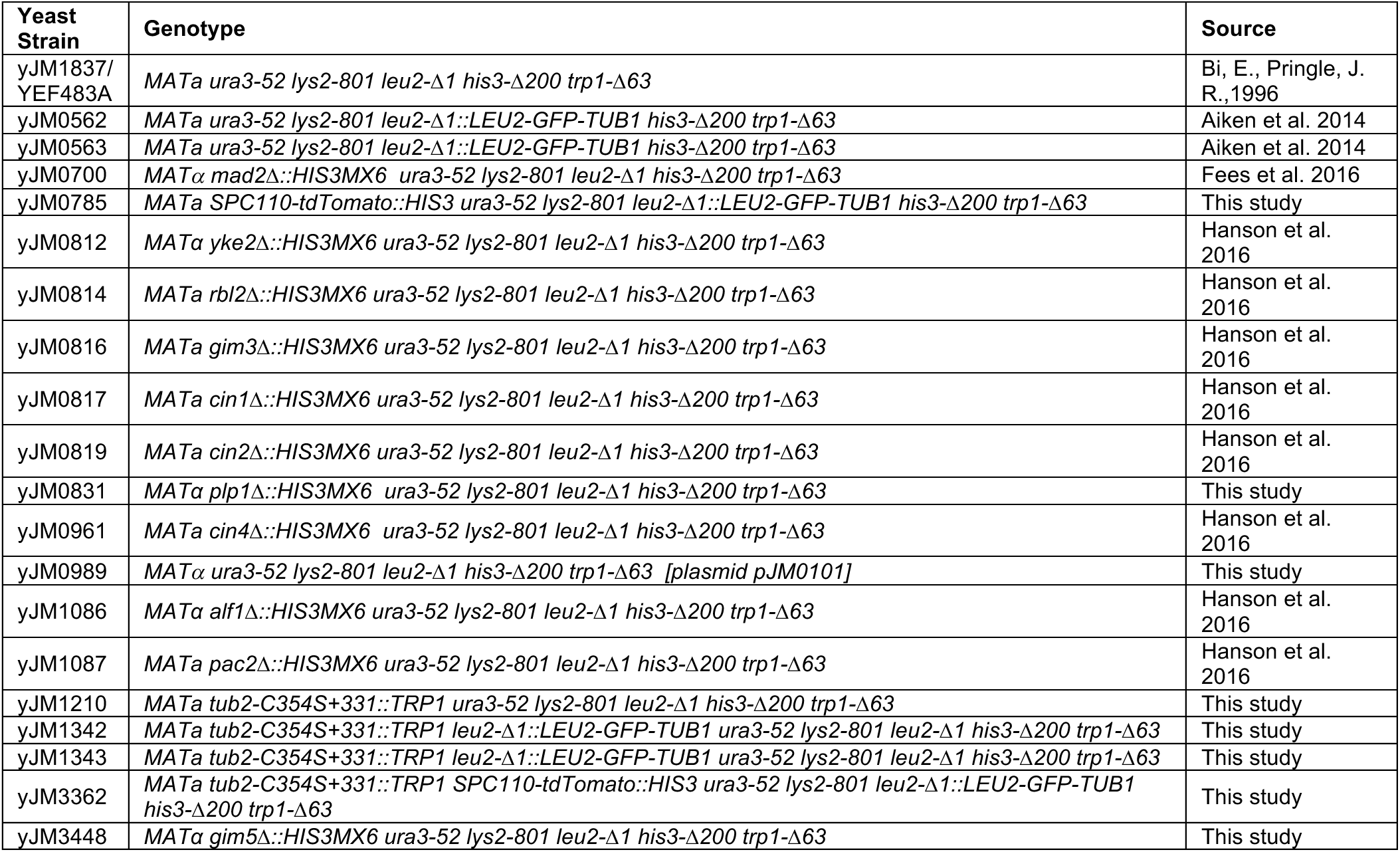
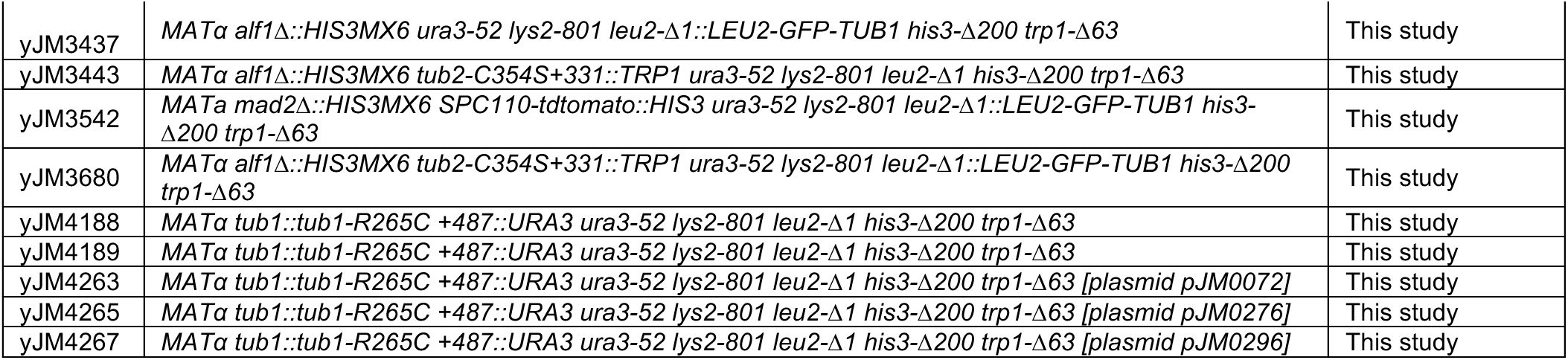
Strains used in this study

