## Supplementary figures and images for "Tubulin recycling limits cold tolerance"

### Supplemental Figure 1

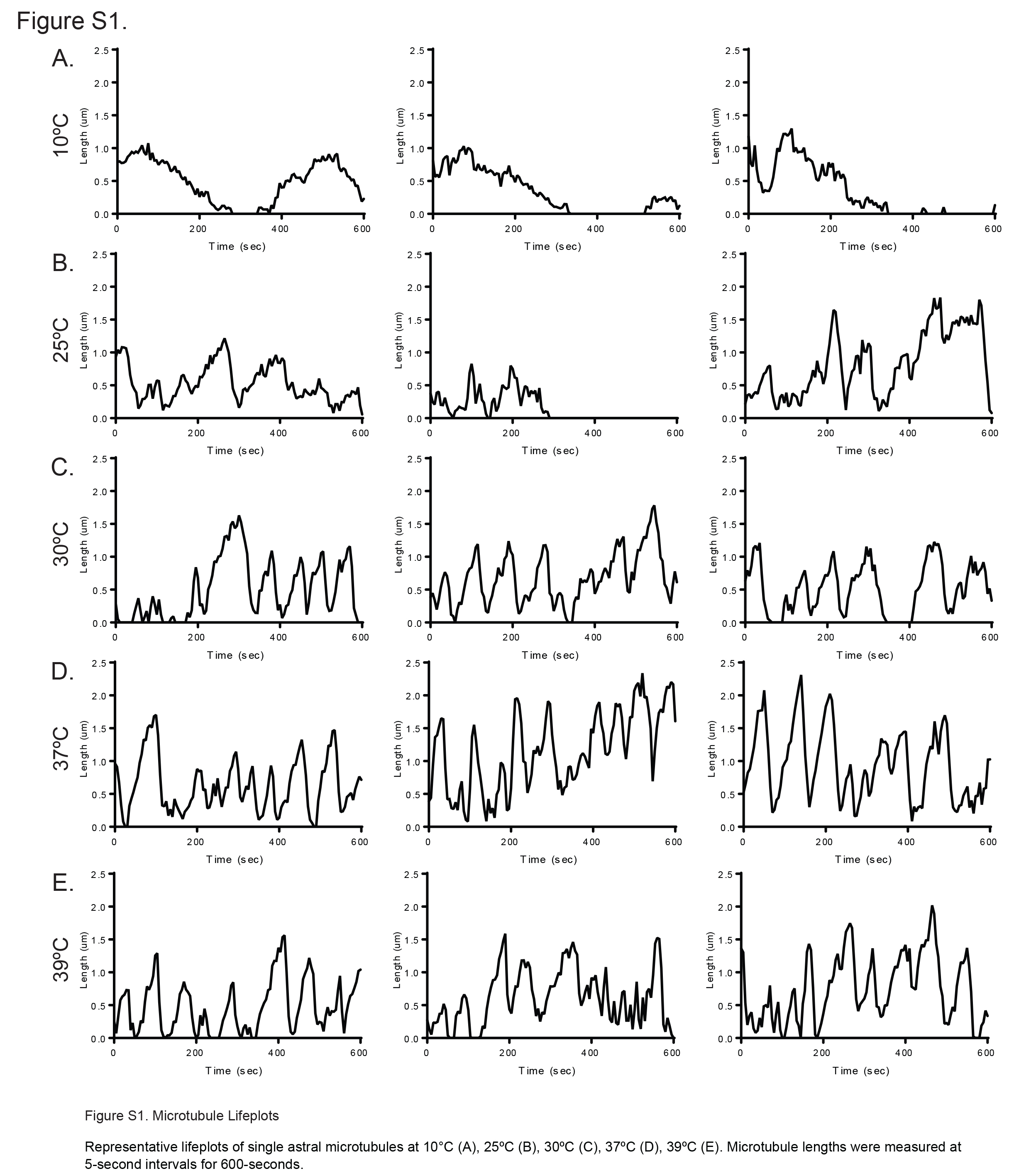

### Supplemental Figure 2

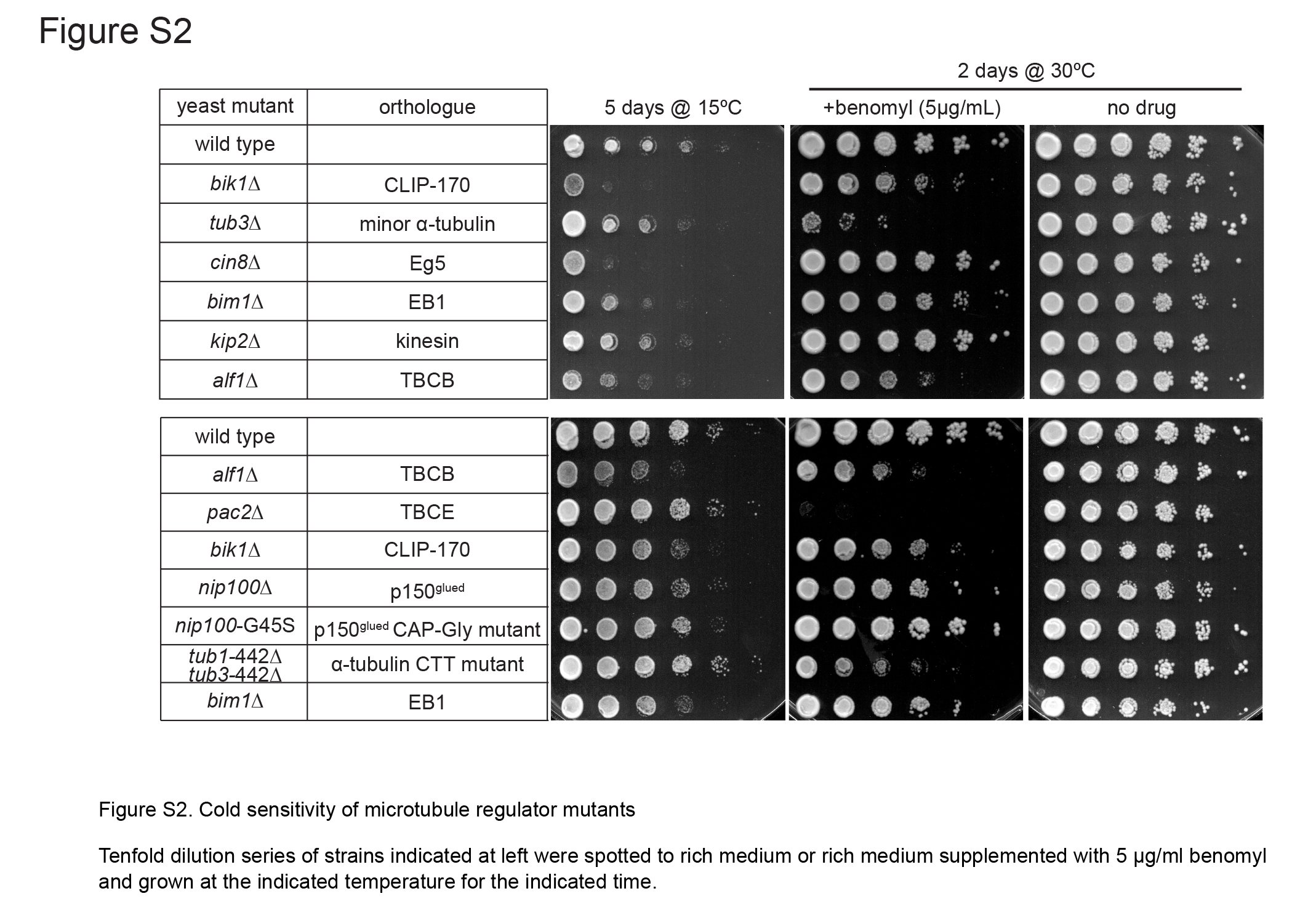

### Supplemental Figure 3

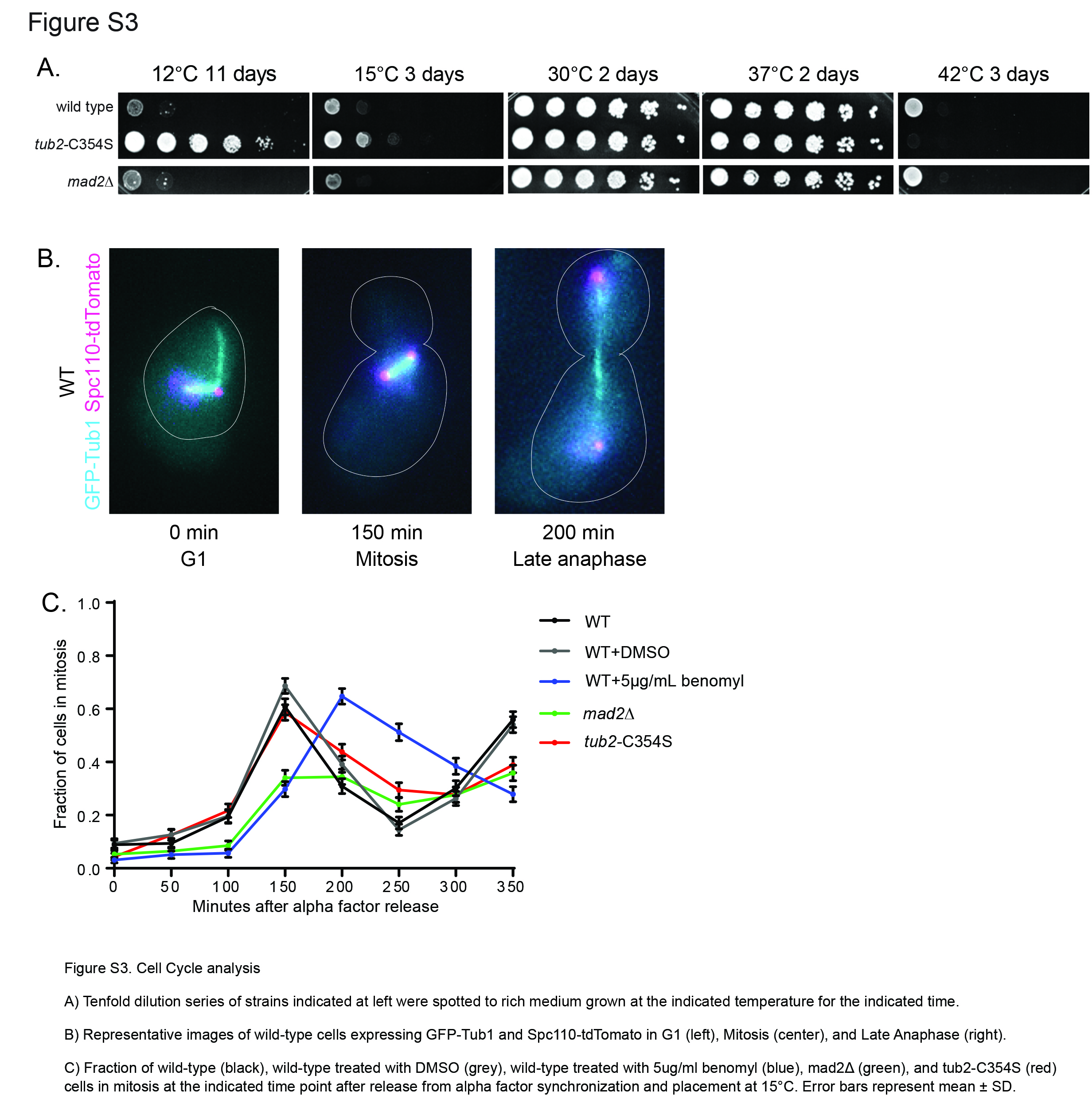
